# Age-related compensatory reconfiguration of PFC connections during episodic memory retrieval

**DOI:** 10.1101/858357

**Authors:** Lifu Deng, Mathew L. Stanley, Zachary A. Monge, Erik A. Wing, Benjamin R. Geib, Simon W. Davis, Roberto Cabeza

## Abstract

During demanding cognitive tasks, older adults (OAs) frequently show greater prefrontal cortex (PFC) activity than younger adults (YAs). This age-related PFC activity increase is often associated with enhanced cognitive performance, suggesting functional compensation. However, the brain is a complex network of interconnected regions, and it is unclear how network connectivity of PFC regions differs for OAs vs. YAs. To investigate this, we examined the age-related difference in functional brain network mediating episodic memory retrieval. YAs and OAs participants encoded and then recalled visual scenes, and age-related differences in network topology during memory retrieval were investigated as a function of memory performance. We measured both quantitative changes in functional integration and qualitative reconfiguration in connectivity patterns. The study yielded three main findings. First, PFC regions were more functionally integrated with the rest of the brain network in OAs. Critically, this age-related increase in PFC integration was associated with better retrieval performance. Second, PFC regions showed stronger performance-related reconfiguration of connectivity patterns in OAs. Finally, the magnitude of PFC reconfiguration increases in OAs tracked reconfiguration reductions in the medial temporal lobe (MTL) – a core episodic memory region, suggesting that PFC connectivity in OAs may be compensating for MTL deficits.

## Introduction

Despite substantial anatomical and functional decline, the aging brain retains a surprising degree of neural plasticity and functional flexibility (Park and Reuter-Lorenz 2009; Cabeza, Albert, et al. 2018). In functional neuroimaging studies, for example, compared to younger adults (YAs), older adults (OAs) tend to exhibit increased neural activity in the prefrontal cortex (PFC) in a wide range of tasks (Cabeza and Dennis 2012; Turner and Spreng 2012; Lighthall et al. 2014; Spreng and Turner 2019). This age-related increase in PFC activation is often attributed to *functional compensation*, which has been defined as a cognition-enhancing over-recruitment of neural resources in response to cognitive demands (Cabeza, Albert, et al. 2018). Most previous studies reporting compensation in OAs have focused on the activity of individual regions without addressing how those regions are interconnected with other brain areas. The human brain is, however, a large-scale complex network of interconnected and interdependent regions, and the transient, dynamic patterns of functional connections between them is essential for human cognition (Bressler and Menon 2010; Medaglia et al. 2015; Cabeza, Stanley, et al. 2018; Stanley et al. 2019). For this reason, age-related compensation can only be successful if it involves not only changes in the activity of individual brain regions but also quantitative and qualitative changes in functional connections between these regions and the rest of the brain. Thus, it is critical to show that the PFC regions over-recruited by OAs change their topological properties within the whole-brain network to facilitate functional compensation.

To investigate this issue, we assessed large-scale functional brain networks in YAs and OAs during episodic memory retrieval. In particular, we had three specific goals. The first goal was to investigate whether PFC regions become more integrated, or functionally interconnected, with the rest of the functional brain network in OAs relative to YAs, and whether this age-related increase in PFC-brain connectivity is associated with better cognitive performance. If a systematic shift in PFC functional connectivity is associated with better cognitive performance in OAs, such result would meet the first of two essential criteria for identifying functional compensation (Cabeza, Albert, et al. 2018). Finding a positive association between network integration and cognitive performance is crucial for interpreting changes as compensatory, because network integration increases are not always advantageous. Some studies using task-related fMRI have linked increased network integration to better cognitive performance in YAs (Westphal et al. 2017; Geib, Stanley, Dennis, et al. 2017) and in OAs (Gallen et al. 2016; Crowell et al. 2019). However, resting-state fMRI studies have linked age-related increases in network integration to a loss of differentiation in the modular network structure (Chan et al. 2014) and a reduction in grey matter volume (Geerligs et al. 2015). Thus, increased PFC integration is not necessarily beneficial for cognitive performance, as it could reflect a reduction in neural specificity. Therefore, a positive association with performance is essential for attributing this effect to functional compensation.

The second goal of current study was to examine whether OAs, relative to YAs, are more likely to *reconfigure* their PFC connections to enhance memory performance. Whereas “integration” refers to quantitative increases in connectivity strength between discrete pairs of cortical nodes in different brain communities or modules (e.g., aging increases connectivity between Regions A, B, and C), “reconfiguration” refers to qualitative changes in the pattern of connections (e.g., aging increases connectivity between Region A and Region B but reduces connectivity between Region A and Region C). That is to say, we focused on whether and to what extent the global connectivity patterns of PFC nodes in the high memory state display dissimilarity with the global connectivity patterns in the low memory state. Converging lines of evidence suggest that transient, dynamic changes in patterns of functional connectivity are crucial for supporting cognitive functions across task domains and demands (Cole et al. 2013; Cole et al. 2014; Spielberg et al. 2015; Simony et al. 2016; Gallen et al. 2016; Geib, Stanley, Dennis, et al. 2017; Geib, Stanley, Wing, et al. 2017; Davis et al. 2018; Cabeza, Stanley, et al. 2018; Stanley et al. 2019). The ability for PFC regions to flexibly reconfigure their functional connections is thought to subserve control-related functions (Cole et al. 2013), including those that aid in memory retrieval (Monge et al. 2018). Therefore, the age-related compensatory role of PFC regions during episodic memory retrieval might be associated with greater reconfiguration of PFC connections.

Finally, the third goal of the study was to investigate whether increased age-related PFC functional reconfiguration is associated with any deficit in another task-related region, such as medial temporal lobe (MTL). The notion of functional compensation implies the existence of a deficit that is being “compensated for.” Thus, it is essential to show that the age-related change in neural activity or connectivity attributed to compensation is associated with an insufficiency in some other neural event thought to support the cognitive function. Such a finding would fulfill the second essential criterion for identifying compensation (Cabeza, Albert, et al. 2018). In healthy YAs, recent research has indicated that increases in memory performance are associated with the reconfiguration of functional connections in the MTL, a region commonly implicated in successful episodic memory retrieval (Geib, Stanley, Dennis, et al. 2017; Geib, Stanley, Wing, et al. 2017). In univariate activation studies, increased PFC activity in OAs has been linked to reduced MTL activity (Hedden et al. 2005; St. Jacques et al. 2009), suggesting that PFC over-recruitment might compensate for MTL deficits in episodic memory (Cabeza and Dennis 2012). It remains unclear, however, whether increased reconfiguration of PFC functional connections is associated with a difficulty of MTL regions to reconfigure their functional connections.

In sum, we investigated if in OAs compared to YAs (1) PFC regions become more integrated with the rest of the network as a function of successful performance (*1st compensation criterion*), (2) PFC show greater performance-related network reconfiguration, and (3) PFC reconfiguration is negatively correlated with MTL reconfiguration (*2nd compensation criterion*). During fMRI scanning, YA and OA participants encoded and then recalled visual scenes with different degrees of success (low vs. high recall). Functional whole-brain networks were constructed from recall trials, with separate *high* and *low* recall networks for each participant (for a similar methodological approach, see (Geib, Stanley, Dennis, et al. 2017; Geib, Stanley, Wing, et al. 2017). Patterns of functional connections were then characterized for *high* and *low* recall networks using several complementary graph theoretic metrics.

## Methods

### Participants

A total of 22 YAs and 22 OAs participated in this study for monetary compensation. Participants were free of significant health problems (including atherosclerosis, neurological and psychiatric disorders), and not taking medications known to affect cognitive function or cerebral blood flow (except antihypertensive agents), according to their self-report. The OAs were further screened with the Beck Depression Inventory (BDI) (Beck et al. 1961) and the modified Mini-Mental State Examination (Folstein et al. 1975) (inclusion criterion ≥ 27, *M* = 29.2, *SD* = 0.7). All participants were right-handed, fluent English speakers, and had completed at least 12 years of education. One YA and one OA were excluded because of missing functional data from the first run caused by a technical error; one additional OA was excluded from analysis due to a poor quality T1 image, which prevented the participant’s functional images to be properly normalized into MNI space. Consequently, these data were analyzed with the remaining 21 YAs (*M_age_* = 23.5 years, *SD* = 3.0 years, age range = [18-30], 9 men, 12 women) and 20 OAs (*M_age_* = 70.5 years, *SD* = 5.4 years, age range = [61-82], 11 men, 9 women). Results from the sample of YAs (but not the sample of OAs) were previously reported in published manuscripts (Geib, Stanley, Wing, et al. 2017; Wing et al. 2015). The Duke University Institutional Review Board approved all experimental procedures, and participants provided informed consent prior to engaging in the experiment.

### Experimental paradigm

Participants completed three encoding runs followed by three retrieval runs **(Fig. 2A)**. During the encoding runs, participants studied pictures of complex scenes (e.g., beach, barn). There were 32 pictures shown in each encoding run, and the order of presentation was randomized within each run across participants. In total, participants viewed 96 pictures of distinct scenes. During each encoding trial (4 sec), participants were presented with a single picture that included a descriptive label below the image (e.g., “tunnel” or “barn”). Participants rated, on a 4-pt scale, the representativeness of the image for the given label (1 = not representative, 4 = highly representative). This task was included to ensure that participants would remain focused and attend to the details of each picture. Each encoding trial was followed by an active baseline interval of 8 sec, during which participants were presented whole numbers ranging from 1 to 4. Participants were instructed to select the button corresponding to the presented number. The retrieval runs were identical in format to the encoding runs with the exception that only the descriptive labels of the encoded pictures were presented. Participants were instructed to recall the scene that previously accompanied the label with as much detail as possible, and rate the amount of memory detail with which they could remember for the specific picture on a 4-pt scale (1 = least amount of detail, 4 = highly detailed memory).

Immediately after the scanning session, participants completed a four-alternative forced-choice recognition test outside of the scanner. For each recognition trial, the target picture from the encoding run and three distractor pictures were presented on the computer screen. Participants selected the picture they believed they saw during encoding. Then, participants rated how confident they were in the preceding recognition decision using a 4-pt scale (1 = guess, 4 = very confident).

### MRI acquisition

MRI data were collected on a General Electric 3T MR750 whole-body 60 cm bore MRI scanner and an 8-channel head coil. The MRI session started with a localizer scan, in which 3-plane (straight axial/coronal/sagittal) localizer faster spin echo (FSE) images were collected. Then, the functional images were acquired using a SENSE spiral-in sequence (repetition time [TR] = 2000 ms, echo time = 30 ms, field of view [FOV] = 24 cm, 34 oblique slices with voxel dimensions of 3.75 × 3.75 × 3.8 mm). The functional images were collected over six runs – three encoding runs and three retrieval runs. There was also a functional resting-state run after the third encoding run, which is not reported here. Stimuli were projected onto a mirror at the back of the scanner bore, and responses were recorded using a four-button fiber-optic response box (Current Designs, Philadelphia, PA, USA). Following, a high-resolution anatomical image (96 axial slices parallel to the AC-PC plane with voxel dimensions of 0.9 × 0.9 × 1.9 mm) was collected. Finally, diffusion-weighted images were collected, which are not reported here. Participants wore earplugs to reduce scanner noise, and foam pads were used to reduce head motion. When necessary, participants wore MRI-compatible lenses to correct vision.

### fMRI preprocessing

For each run, the first six functional images were discarded to allow for scanner equilibrium. All functional images were preprocessed in a SPM12 (London, United Kingdom; http://www.fil.ion.ucl.ac.uk/spm/) pipeline. The functional images were slice time corrected (reference slice = first slice), realigned to the first scan in the first session, and subsequently un-warped. Then, the functional images were co-registered to the skull-stripped high-resolution anatomical image (skull-stripped by segmenting the high-resolution anatomical image and only including the gray matter, white matter, and cerebrospinal fluid segments). The functional images were normalized into MNI space using DARTEL (Ashburner 2007); the study specific high-resolution anatomical image was created using all of the study participants. The voxel size was maintained at 3.75 × 3.75 × 3.8 mm^3^ and the normalized-functional images were not spatially smoothed. Lastly, the DRIFTER toolbox (Särkkä et al. 2012) was used to denoise the functional images.

### Network construction

We created functional brain networks for each participant using a beta time-series analysis (Rissman et al. 2004; Geib, Stanley, Wing, et al. 2017; Geib, Stanley, Dennis, et al. 2017). The nodes in the network were defined by the 90 regions of interest (ROIs) in AAL atlas, excluding cerebellar regions (Tzourio-Mazoyer et al. 2002)^1^. The beta time-series of each node was obtained by averaging the beta-estimate of each trial within that ROI. To obtain the beta-estimates for each trial, we conducted a single-trial model analysis within a general linear model. This approach estimates a first-level model in which one regressor models a specific event of interest and another regressor models all the other events (Mumford et al. 2012). This process was repeated on every trial. Each trial was modeled with a standard hemodynamic response function (HRF) with the temporal and dispersion derivative. Each model also included the six raw motion regressions, a composite motion parameter (derived from the Artifact Detection Tools [ART]^2^), outlier TRs (scan-to-scan motion > 2.0 mm or degrees, scan-to-scan global signal change > 9.0 z-score; derived from ART), the white matter time-series, and cerebrospinal fluid time-series. We also modeled the temporal and dispersion derivatives and implemented a 128 sec cutoff high-pass temporal filter.

Beta-values for each trial were then matched to the memory detail scores reported by participants during each trial on the 4-pt scale. Beta-values obtained during trials where participants reported high levels of detail during recollection (ratings of 3 and 4) were combined into a beta-series for constructing the functional brain networks in the *high* memory condition; beta-values obtained during trials where participants reported low levels of detail during recollection (ratings of 1 and 2) were combined into a beta-series for constructing the functional brain networks in the *low* memory condition. For each memory condition (i.e. *high* vs. *low* memory), a connectivity matrix was constructed for each participant by calculating the Pearson’s correlation of the condition-specific beta-series between all possible pairs of regions.

Because the *high* and *low* memory networks were often constructed with an unequal number of trials, a bootstrap process was employed to eliminate the potential confound that observed differences in network topology could be the product of unequal trial counts in functional network construction. For the condition (i.e., *high* or *low* memory) with more trials, a subset of *N* trials was randomly sampled, where *N* equals the number of trials in the other condition. This process was repeated 500 times, creating 500 connectivity matrices for the bootstrapped condition. Finally, the 500 connectivity matrices were averaged into one representative matrix for the bootstrapped condition. For network analyses, the diagonal values of the obtained connectivity matrix were set to zero.

### Network analyses

#### Modularity

We started the network analysis by identifying functional sub-systems, or modules. In brief, a modularity analysis based on the Louvain algorithm (Blondel et al. 2008; Rubinov and Sporns 2010) was used to detect common modular structure of the retrieval network in order to allow comparison of network measures across groups and condition (i.e., memory performance). Extra steps were included in the analysis to ensure that the modules would not bias specific age group or condition, which are detailed as follows. In the first step, a *common representative network* was created in a two-step manner. We first averaged the connectivity matrices for each age group (YA/OA) in each memory condition (High/Low) individually, and then we averaged these four age and condition-specific networks. No significant difference in the average connectivity level was found between YAs and OAs (t(39)= −0.73, p= 0.47), which ensured that the common representative network was not biased by either age group. Then, in the second step, a mask of strong connections was created by obtaining a set of X% (X tested in a range between 5 and 30 with an increment of 1) strongest connections for each *age-specific representative network* (averaged across condition), and then performing union operation on these two sets of strong average connections. The mask was then used to threshold the *common representative network* (averaged across age and condition), setting the weak connections to zeros. Here, the rationale for using a thresholded, weighted network for module detection instead of a binary network or a fully weighted network (i.e., no threshold at all) is that it allows us to optimally utilize the information in strong connections and, at the same time, exclude weak or negative connections that are difficult to interpret. Finally, the thresholded, weighted common representative network was submitted to the Louvain algorithm using a default resolution parameter (gamma) of 1. The algorithm was run 1000 times on this network, and the optimal module assignment with the highest modularity *Q* value was elected as the optimal module assignment under the given sparsity.

The similarity between module assignments across different levels of sparsity was quantified using a partition-similarity algorithm based on mutual-information (Meilă 2007). Across the different sparsity levels, we consistently obtained a 6-module partition (see **Fig. 1**) when the sparsity of age-specific network was between 5% and 16%, and when the sparsity was between 5% and 10%, the module assignments were almost identical **(Fig. S1a)**. We used the module assignment at 10% sparsity as the module definition in our following analysis, because (i) this partition is clearly comparable to existing research on memory networks in aging (Gallen et al. 2016; Monge et al. 2018), (ii) it has the highest average consensus with the partition results at other thresholds (**Fig. S1b**), and (iii) this sparsity level ensures that the network will not fracture, allowing us to appropriately investigate connectivity between different modules. In addition, we computed the similarity of common module definition to the individualized module partition for each participant and compared this similarity measure between the two groups. The module partitions in OAs and YAs showed no difference in similarity to the common module definition (t(39)=1.13, p=0.27), confirming that the common module assignment was not significantly biased by one of the two age groups.

**Figure 1.**
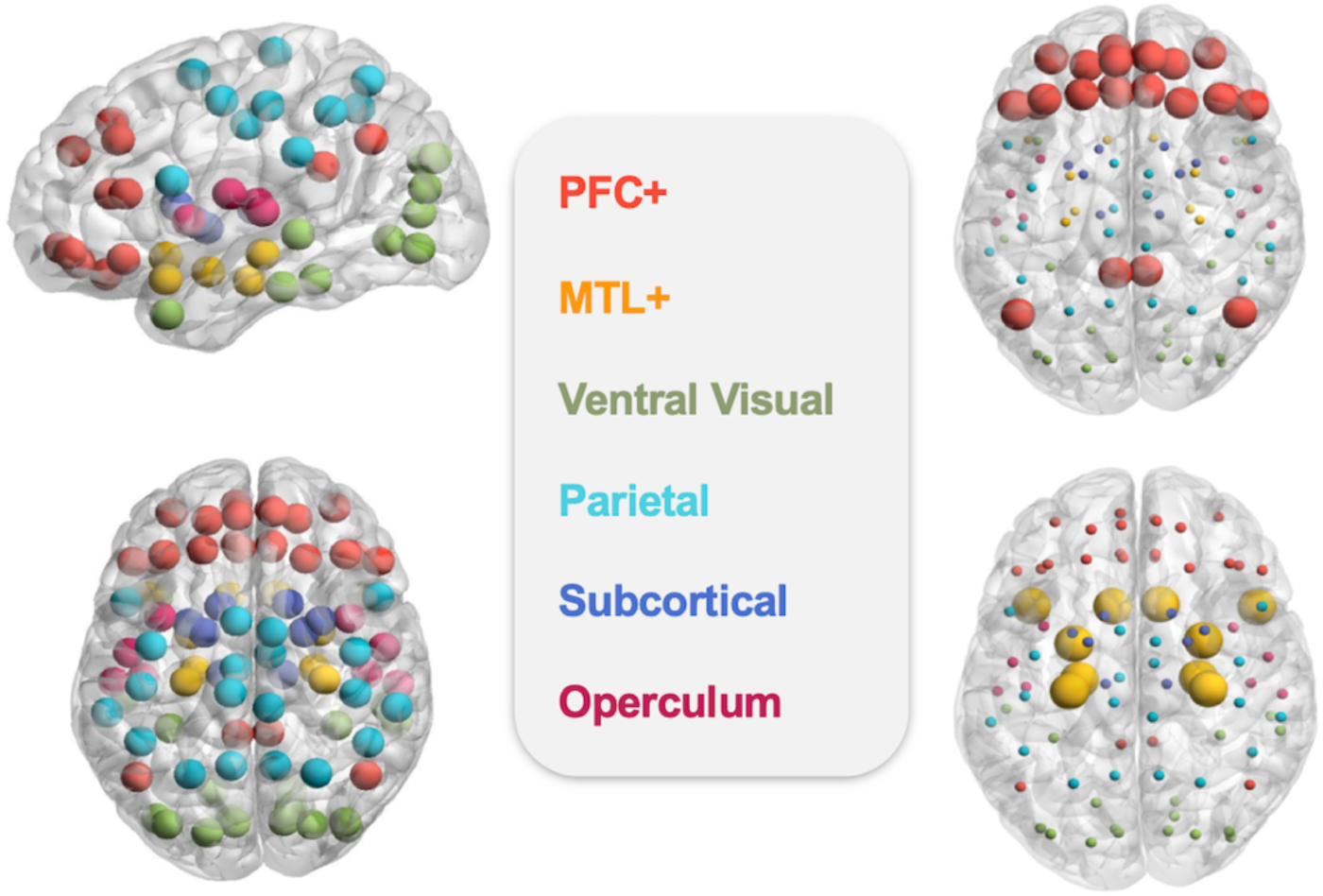
Modularity analysis revealed six modules in the common task network, including the PFC+ module (top right) and MTL+ module (bottom right).

**Figure 2.**
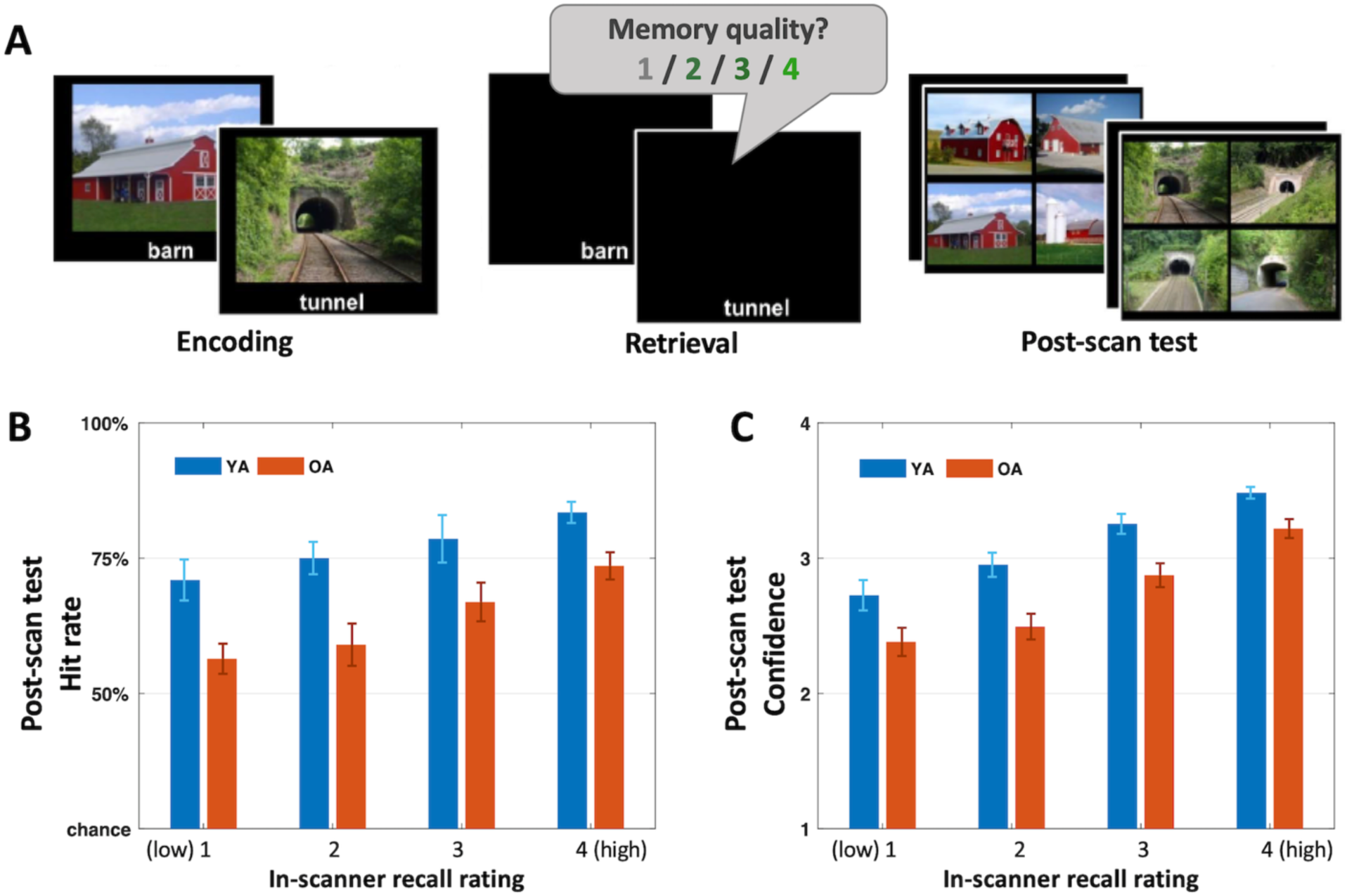
**A)** Experimental paradigm. **B)** Post-scan recognition hit rate as a function of in-scanner recall rating in YAs and OAs. **C)** Post-scan recognition confidence as a function of in-scanner recall rating in YAs and OAs. Error-bars indicate standard error of the mean (SEM).

#### Module connectivity properties

To quantify the integrative properties of the obtained modules, we computed within- and between-module connectivity (Guimerà and Amaral 2005). The within-module functional connectivity refers to the average functional connectivity between all pairs of regions contained within the same module; the between-module functional connectivity refers to the average of all functional connections linking the given module to other modules. Finally, a composite measure of whole-brain integration for each module was defined as the ratio of between-module functional connectivity to within-module functional connectivity:

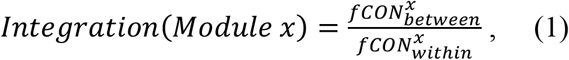

This metric reflects the tendency of a specific module to exhibit functionally interactions with other modules in the larger network. For the convenience of illustration, in the *Results* section the above analyses are based on 10% network sparsity (age-specific network), which is consistent with the sparsity used in the modularity analysis. It is worth noting that the results remained highly consistent across different network sparsity levels. These results are available in the Supplementary Materials (**Figs. S2 & S3**).

#### Reconfiguration

We defined the reconfiguration of functional connections between *high* memory and *low* memory conditions as one minus the Spearman’s correlation of one node’s connectivity vectors in *high* and *low* memory conditions (see **Fig. 5A**):

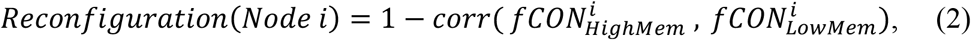

To fully characterize the connectivity pattern and equalize the vector length for all nodes, the input connectivity vectors are based on non-thresholded network, containing all connectivity values. In consistency with previous studies (Geib, Stanley, Wing, et al. 2017; Geib, Stanley, Dennis, et al. 2017), the reconfiguration values within each subject were *z*-scored before submitting to statistical analyses to control for between-subject variability. To ensure that fCON reconfiguration was not simply induced by statistical noise, we performed a bootstrap analysis, where the memory ratings for all trials were randomly shuffled to create surrogate *high* and *low* memory conditions **(Fig. 5B)**. Bootstrapped fCON reconfiguration was calculated from the surrogate *high* and low memory fCON matrices. This process was repeated 1000 times for each subject in order to approximate a comparable null distribution of fCON reconfiguration that was not related to memory condition.

Statistical testing was carried out using R and the statistical toolbox in MATLAB (The MathWorks, Inc.). Participants’ behavioral data and fCON network metrics were analyzed with repeated measures ANOVAs. The brain networks were illustrated with the BrainNet Viewer (http://www.nitrc.org/projects/bnv/)(Xia et al. 2013).

## Results

### Behavioral results

During recall in the scanner, participants rated the quality of their memory of the images from 1 (*least amount of detail*) to 4 (*highly detailed memory*) **(Fig. 2A**, middle). Outside the scanner, they performed a memory recognition task in which they attempted to identify the encoded scene among 3 similar distractor scenes, and then they rated their confidence **(Fig. 2A**, right). To confirm the objective validity of subjective in-scanner ratings, we analyzed recognition performance outside the scanner as a function of in-scanner ratings. We performed separate 2 (age: YA vs. OA) x 4 (in-scanner recall rating: 1-4) ANOVAs for accuracy and confidence as separate outcome variables. In these ANOVAs, higher in-scanner recall ratings were associated with higher post-scan recognition accuracy (**Fig. 2B**; F(3,117)=13.17, p=1.8e-7) and confidence **(Fig. 2C;** F(3,117)=64.62, p=2e-16), confirming that in-scanner memory rating reliably tracked memory performance, In the ANOVAs, there were also significant main effects of age on accuracy (F(1,39)=14.73, p=4.4e-4) and on confidence (F(1,39)=14.99, p=4.0e-4), consistent with age-related episodic memory decline. Neither ANOVA yielded reliable interactions. In summary, the behavioral results confirmed the validity of in-scanner recall ratings, which we use to split retrieval trials to create separate networks for High vs. Low Recall.

### Network analyses

The following network analyses had three main goals: (1) investigate if PFC regions become more integrated with the rest of the brain in OAs than YAs, as a function of cognitive performance (first criterion of compensation); (2) examine if age-related changes in PFC recruitment involve a *qualitative* change in performance-related reconfiguration of connectivity patterns; and (3) investigate if PFC reconfiguration is associated with decreased MTL reconfiguration (second criterion of compensation). As noted before, our modularity analysis on the combined YA-OA connectivity data yielded 6 modules (see **Fig. 1**). Our goals focus on two modules of interest, PFC and MTL. The PFC+ module primarily contained PFC regions, but it also included bilateral posterior cingulate cortices and angular gyri. The MTL+ module contained mainly MTL regions, such as bilateral hippocampi, para-hippocampus cortices, and amygdala, but it also included the temporal poles and olfactory cortex.

#### Goal 1: PFC integration and the link to cognitive performance

##### PFC integration

Whole-brain integration (i.e., the ratio of between-module fCON to within-module fCON) was calculated for each module from each individual’s fCON network (thresholded weighted network, including both high and low memory trials). Although the focus of our integration analyses was the PFC+ module, for sake of completeness we also investigated the effects on aging on all modules. An Age-by-Module repeated-measures ANOVA was performed, with module-specific whole-brain integration as the dependent variable. This analysis yielded a significant main effect of Module (F(5,195)=17.11, p=5.13e-14). Indeed, the two modules of interest (PFC+ and MTL+) showed higher module-specific whole-brain integration when compared with the average integration of the other 4 modules (paired t-test within age groups; PFC+ vs. other 4 modules: t_YA_=2.36, p_YA_=0.028; t_OA_=4.38, p_OA_=3.22e-4; MTL+ vs. other 4 modules: t_YA_=5.52, p_YA_=2.07e-5; t_OA_=1.96, p_OA_=0.064; FDR uncorrected), presumably reflecting the contribution of these regions to integrative processing in the network. Importantly, the ANOVA yielded significant Age-by-Module interaction (F(5,195)=13.08, p=5.52e-11), indicating that aging had differential impact on whole-brain integration for different modules.

To further examine the age-related differences on the two modules of interest, we repeated our analysis on only the PFC+ and MTL+ module. Indeed, we again observed a significant Age-by-Module interaction (F(1,39)=13.44, p=7.32e-4) due to different age effect on integration for PFC+ and MTL+ modules (see **Fig. 3**). In PFC+ module, integration with the rest of the brain network was significantly greater in OAs than YAs (t(39)=2.26, p=0.029), whereas in the MTL+ module there was a significant age difference in the opposite direction (t(39)=2.65, p=0.011). In sum, aging was associated with increased whole-brain integration in the PFC+ module and decreased whole-brain integration in the MTL+ module. This opposite effect of aging on PFC+ and MTL+ module organization is further examined below in reconfiguration analyses on differential effects of aging on PFC+ vs. MTL+ reconfiguration (*Goal 2*).

**Figure 3.**
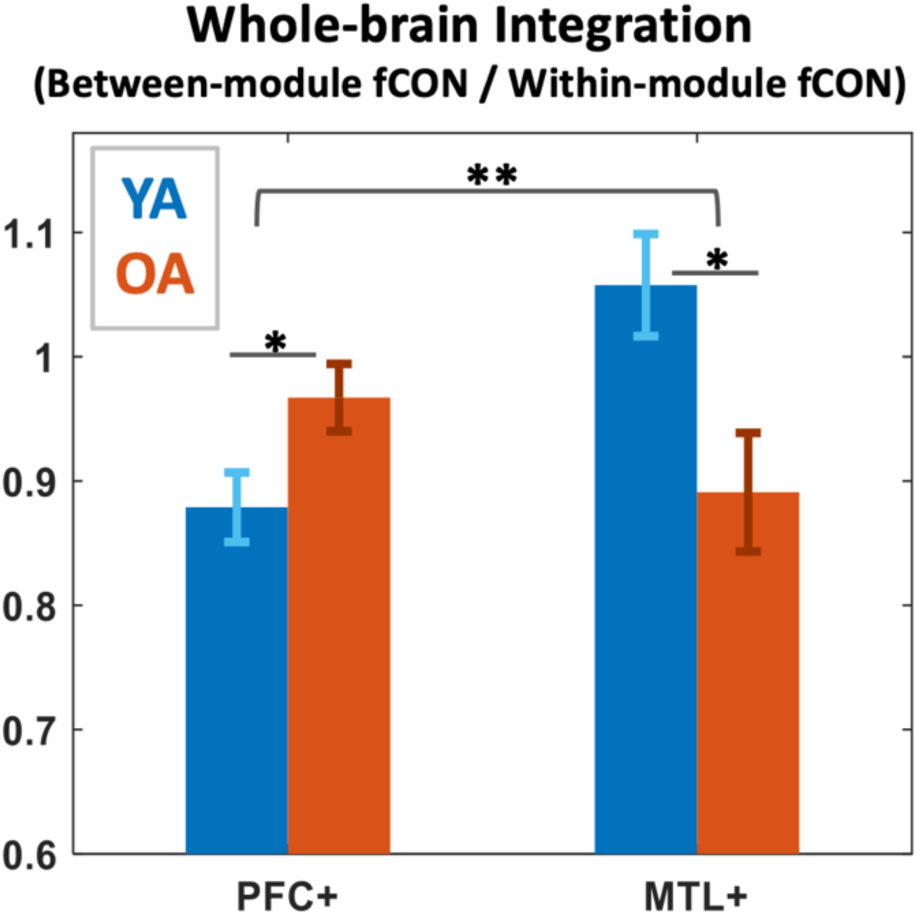
Compared with YAs, OAs showed increased network integration in PFC regions. Horizontal bracket indicates Age x Module effect; horizontal line indicates two-sample t-test (**: p<0.01; *: p<0.05). Error-bars indicate SEM.

##### Link to cognitive performance

We then investigated whether age-related increase of PFC’s role in the network contributes to cognitive performance in OAs. We split the data into high/low memory trials and examined if the integration measure of the PFC+ module varied with memory success; this analysis did not yield reliable reliable Memory effect (F(1,39)=0.18, p=0.67) or Age-by-Memory effect (F(1,39)=0.82, p=0.37). Given that the integration is the ratio of between-module fCON to within-module fCON, we then investigated age-related difference in each of these two types of fCON. We performed 2 (Age group: YA vs. OA) x 2 (Memory: high vs. low) ANOVA on within-module and between-module connections separately. For within-module fCON (**Fig. 4A**), the ANOVA yielded a significant main effect of Age (F(1,39)=4.92, p=0.033), but no reliable main effect of Memory (F(1,39)=2.27, p=0.14), or Age-by-Memory interaction (F(1,39)=1.38, p=0.25). In contrast, for between-module fCON, the ANOVA yielded no reliable main effect of Age or Memory, but a significant Age-by-Memory interaction (F(1,39)=5.09, p=0.030). As illustrated by **Fig. 4B**, this interaction reflected the fact that fCON connecting PFC+ module to other systems in the brain was greater for high than low memory trials in OAs (t(19)=2.52, p=0.021, FDR corrected) but not in YAs (t(20)=−0.66, p=0.51). This finding provides evidence for a significant link between increased PFC connectivity in OAs and successful memory performance (*1^st^ compensation criterion*).

**Figure 4.**
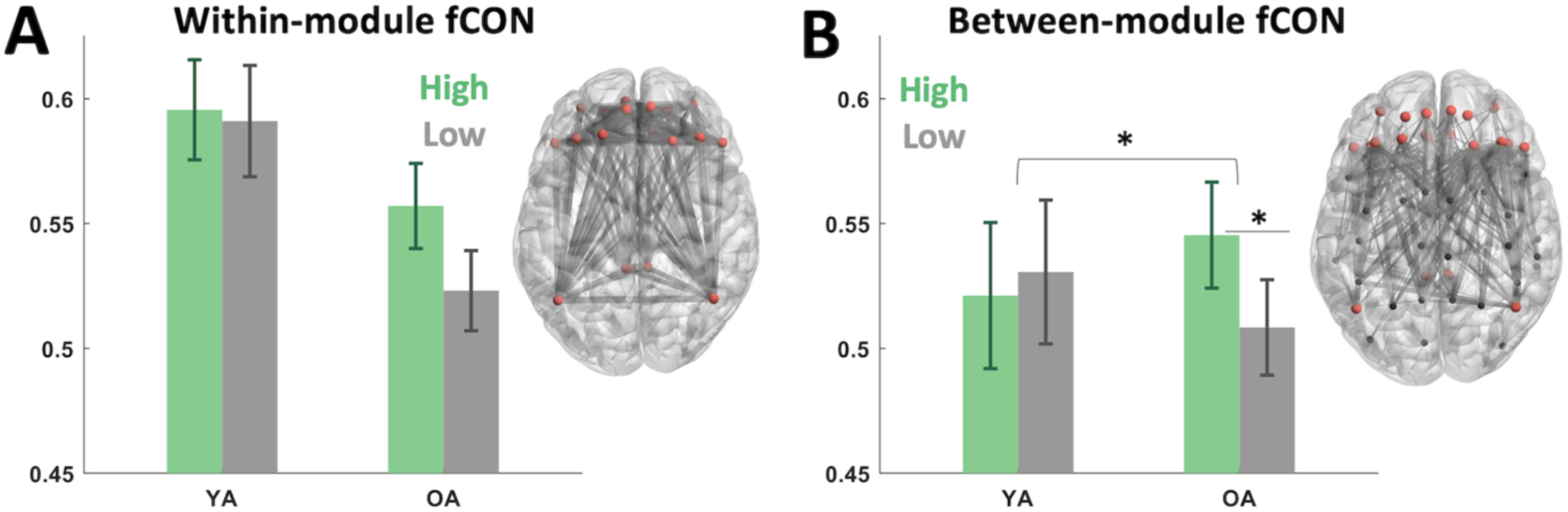
**A)** Average fCON within PFC+ module. **B)** Average fCON between PFC+ module and all other modules. Red nodes indicate brain regions in PFC+ module, while black nodes indicate regions belonging to other modules. Horizontal bracket indicated a significant Age x Module interaction; underscores indicated paired t-test (*: p<0.05). Error-bars indicate SEM.

**Figure 5.**
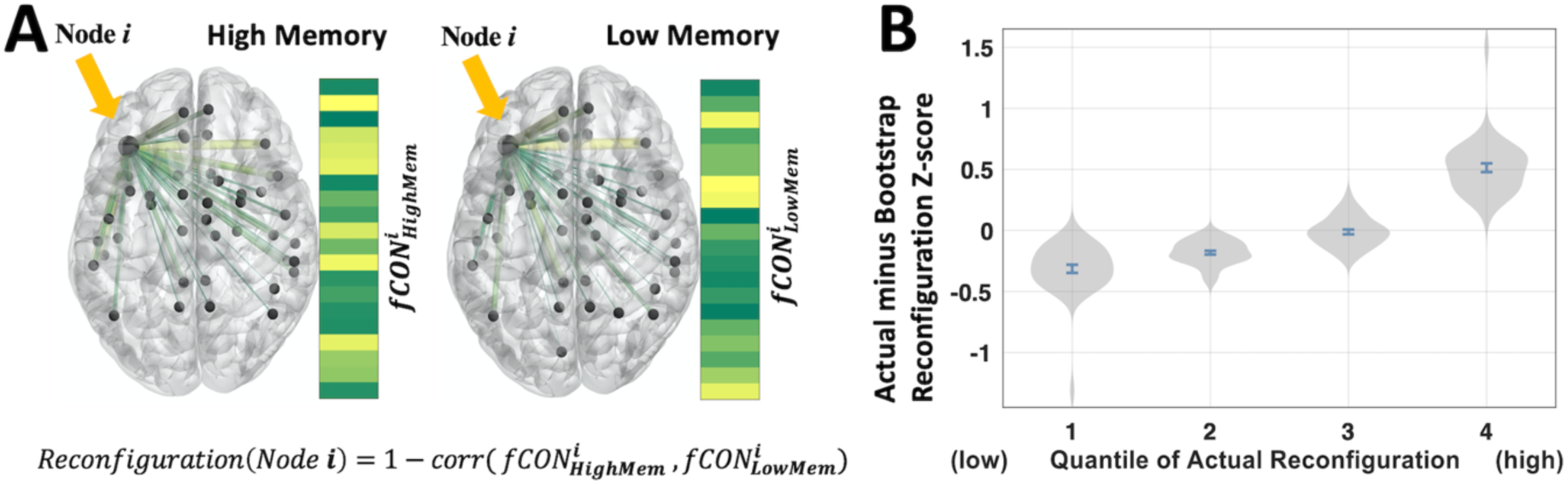
A) An illustration of the computation of memory-related reconfiguration. **B)** The differences between actual and bootstrapped reconfiguration were binned into 4 equal-size samples for each subject according to the actual reconfiguration value.

#### Goal 2: Aging and reconfiguration of PFC connectivity

The second main goal of the study was to examine if age-related differences in PFC recruitment involve a change in network reconfiguration. The change of fCON between different memory states is multivariate in nature; thus changes in fCON may not be fully observable by measurements such as average fCON strength. Therefore, the fCON reconfiguration between memory conditions was measured in each ROI for each subject **(Fig. 5A)**. As illustrated in **Figure 6**, PFC+ reconfiguration from low to high memory was greater for OAs than YAs, suggesting that the effects on aging on PFC connectivity are not merely quantitative; OAs qualitative reorganize the PFC connections as a function of memory performance.

**Figure 6.**
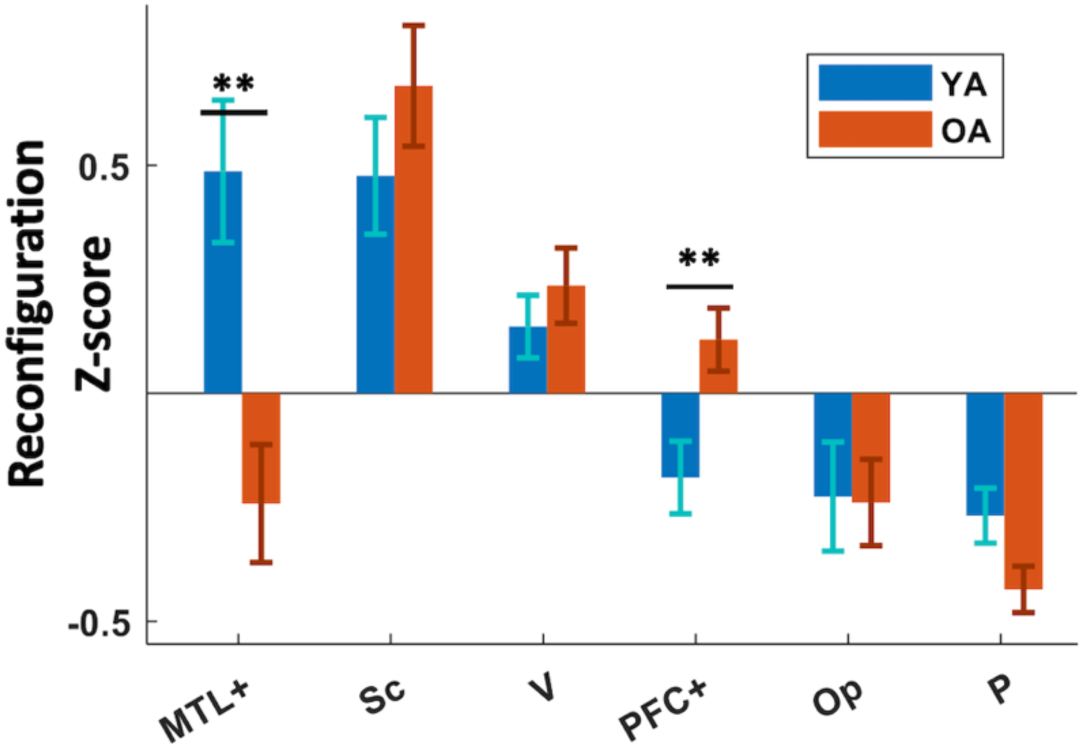
The value distribution of reconfiguration across all modules. Error-bars indicate SEM. (**: p<0.01)

One premise is required for further analysis on connectivity reconfiguration: that this measure should reflect real memory-related change in connectivity pattern, rather than simply be driven by the instability of connectivity computation due to sub-sampling the total set of trials (i.e., drawing the high and low memory trials from the trial set and computing the connectivity based on each drawing). Thus, we compared the actual reconfiguration values to random reconfiguration by subtracting the average bootstrap reconfiguration values from the actual reconfiguration values for each node in each participant. We then examined the actual-random difference as a function of the magnitude of the actual reconfiguration values **(Fig. 5B)**. Indeed, at the highest quantile, actual reconfiguration values were significantly greater than the expected random reconfiguration (t(40)=14.58, p= 1.35e-17), whereas lower actual reconfiguration values in the lowest two quantiles are significantly below the expected random reconfiguration (first quantile: t(40)= −9.47, p= 9.00e-12; second quantile: t(40)= −11.90, p= 1.03e-14). Such difference in distribution indicates that the reconfiguration measure captures memory-related, multivariate change in fCON network that is substantially different from the statistical null distribution, therefore securing the validity of the following analyses into this measure.

#### Goal 3: Relationship between age effects on PFC vs. MTL reconfiguration

Our third goal was to investigate if PFC reconfiguration is negatively correlated with MTL reconfiguration. To examine this idea, we started by measuring reconfiguration in YAs and OAs in all six network modules **(Fig. 6)**. An 2 (Age group) x 6 (modules) ANOVA yielded a significant Age x Module interaction (F(5,195)= 5.59, p= 7.7e-5), and *post hoc* tests showed that the only modules showing significant age-related differences in integration were PFC+ and MTL+ (t_PFC+_(39)= −2.85, p_PFC+_= 0.0069; t_MTL+_(39)= 3.57, p_MTL+_= 0.00095, FDR corrected).

Given that age-related differences in PFC+ and MTL+ modules were in opposite directions, we performed another ANOVA included just these two modules to investigate the interaction (**Fig. 7A**). The ANOVA yielded a significant Age x Module interaction (F(1,39)=14.71, p=4.5e-4), because YAs showed stronger reconfiguration in MTL+ than PFC+ (t_YA_(20)=3.17, p_YA_=0.0048), whereas OAs displayed the opposite pattern (t_OA_(19)=2.22, p_OA_=0.039).

**Figure 7.**
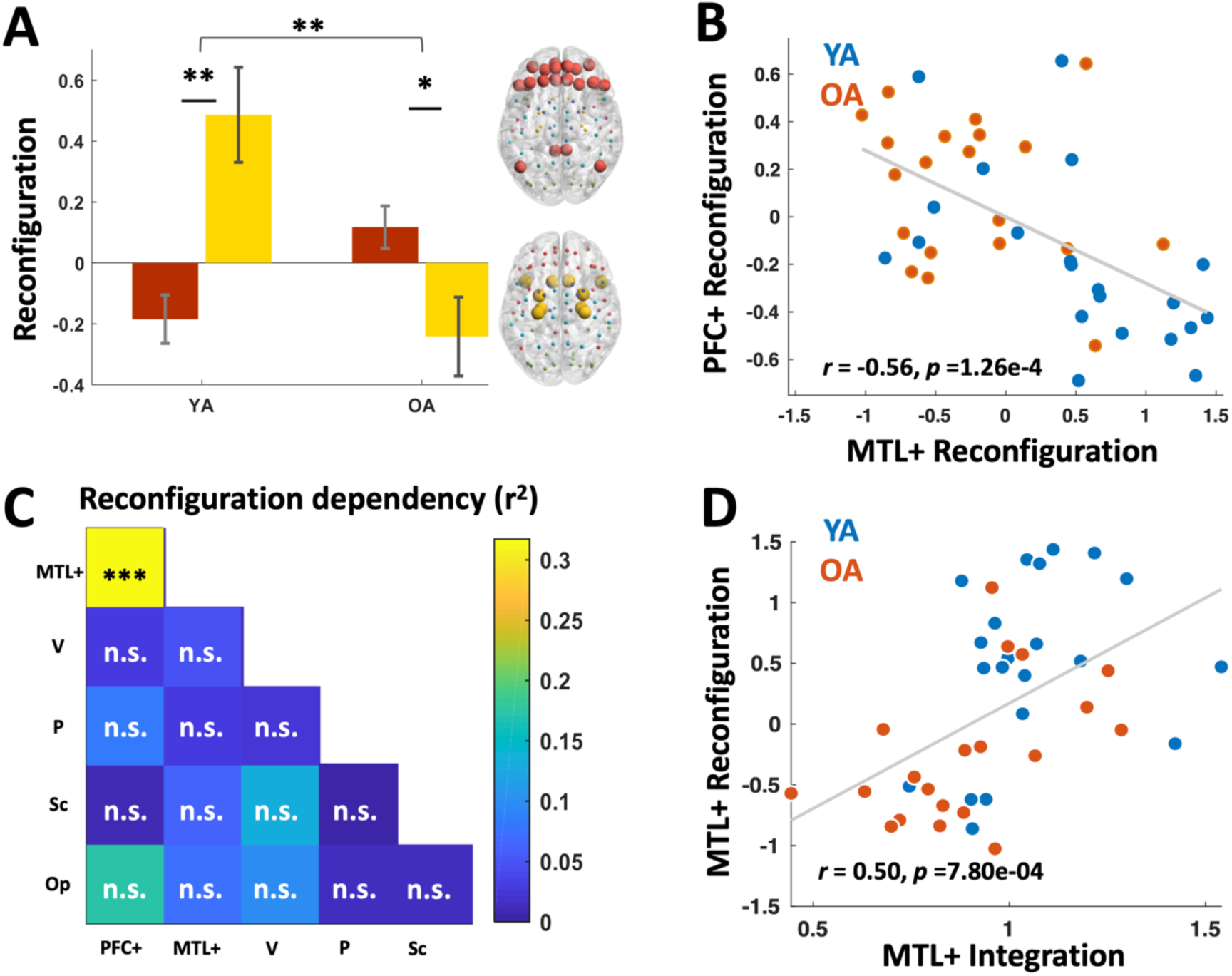
**A)** The values of reconfiguration in PFC+ and MTL+ modules. Horizontal bracket indicated a significant Age x Module interaction; underscores indicated paired t-test (*: p<0.05; **: p<0.01). Error-bars indicate SEM. **B)** The dependency between reconfiguration in MTL+ and PFC+ modules across all subjects. **C)** The dependency (Pearson’s r-square) of reconfiguration between all possible combination of modules (***: p<0.001, FDR corrected; n.s.: not significant). **D)** The dependency between reconfiguration and integration in MTL across all subjects

The results in **Figure 7A** suggest that PFC reconfiguration in OAs compensates for reconfiguration deficit in MTL (2^nd^ criterion of compensation). To investigate this hypothesis, we correlated reconfiguration in these two brain regions. As illustrated by **Fig. 7B**, consistent with compensation, we found a negative correlation between PFC+ and MTL+ reconfiguration across all subjects (Pearson’s r=-0.56, p=1.26e-4). Importantly, across all possible pairs of modules, this significant anti-correlation was present uniquely between MTL+ and PFC+ modules **(Fig. 7C)**. This negative dependency between PFC+ and MTL+ reconfiguration remained in an additional partial correlation analysis, where the age group was regressed out as the variable being controlled (r=-0.45, p=0.0034).

Furthermore, if reduced MTL reconfiguration were indicative of a deficit, this should be convergently supported by other network measures. To explore this possibility, we conducted a *post-hoc* correlation analysis between MTL reconfiguration and integration **(Fig. 7D)**. A significant positive correlation between integration and reconfiguration in MTL (Pearson’s r=0.50, p=7.80e-4) indicated that, in OAs, the reduced MTL reconfiguration is accompanied by a weakened MTL integration into global brain network. This reconfiguration-integration relationship exists only in the MTL module after correcting for multiple comparisons. In summary, fulfilling the 2^nd^ compensation criterion, we observed a negative association with increased PFC reconfiguration in OAs and reduced MTL reconfiguration.

## Discussion

A fundamental goal in the cognitive neuroscience of aging is to understand the variability of age-related cognitive decline across individuals, especially when that decline negatively impacts their everyday lives (Salthouse 2004; Habib et al. 2007; Cabeza, Albert, et al. 2018). The notion of *functional compensation* in aging offers not only a theoretical foundation to better understand differences in cognitive abilities across older adults, but also valuable information regarding possible interventions even after neural degeneration occurs. By leveraging recent advances in network neuroscience, our study investigates how certain brain regions, through their patterns of functional connections, play a different role within the large-scale brain networks of OAs (relative to YAs) to elicit cognition-enhancing effects attributable to functional compensation.

This study yielded three main findings. First, PFC regions became more functionally integrated with the rest of the brain network in OAs relative to YAs, and critically, the increased connectivity from PFC regions was associated with improved memory performance within OA individuals. Second, PFC regions tended to reconfigure their functional connections to support better memory performance in OAs. Third, the extent of functional reconfiguration of PFC connections in aging closely tracked the extent of reconfiguration deficits of MTL regions thought to play a critical role in successful memory retrieval. Taken together, our findings provide strong support for the age-related functional compensation of PFC regions during episodic memory retrieval. By satisfying these essential requirements for identifying cases of functional compensation (Cabeza, Albert, et al. 2018), our findings offer the first evidence for age-related functional compensation at the level of large-scale networks. We discuss each of the three main findings below.

### Retrieval performance in OA supported by PFC integration

Our first goal was to investigate whether PFC regions become more functionally interconnected with the rest of the functional brain network in OAs, and whether this is associated with better cognitive performance. Theorists have suggested that through increased activation of PFC regions, some OAs are able to maintain relatively high cognitive functioning during demanding tasks—including episodic memory tasks—when MTL or occipital cortical function is impaired (Grady et al. 1995; Davis et al. 2008; Park and Reuter-Lorenz 2009; Cabeza and Dennis 2012; although, see Morcom and Henson 2018). These PFC regions are thought to serve control and monitoring functions that aid in the successful retrieval of information from memory (Cabeza and Dennis 2012; Eichenbaum 2017; Inman et al. 2018).

While these previous studies have provided valuable information of *what* regions play compensatory roles in aging, recent developments in network neuroscience may bring further insights into *how* compensation is achieved. There has been growing consensus that although many cognitive functions can be attributed to specific ‘core’ brain regions, they are in fact carried out collaboratively by many different regions (Bressler and Menon 2010; Sporns 2014; Medaglia et al. 2015; Stanley et al. 2019). Episodic memory retrieval, for example, is associated with the activation of many disparate brain regions, which play different but complementary roles, such as storing and indexing memory traces, and evaluating the quality of retrieval (Cabeza and Nyberg 2000; Wagner et al. 2005; Cabeza 2008; Spaniol et al. 2009; Huijbers et al. 2011; Rugg and Vilberg 2013), and is also supported by increased connectivity across the network (Geib, Stanley, Dennis, et al. 2017; Geib, Stanley, Wing, et al. 2017; Westphal et al. 2017). Thus, our results are consistent with the heuristic that if the PFC regions do play a compensatory role in the memory retrieval process, then this role should be realized through network-level interactions.

To be compensatory, a brain region might exhibit increased direct and indirect communication with other task-related regions, leading to a highly integrative network architecture. Indeed, among resting-state fMRI studies, greater integration between disparate network modules during normal aging has been consistently observed (e.g., Betzel et al. 2014; Chan et al. 2014; Geerligs et al. 2015; Grady et al. 2016). However, resting-state studies cannot fully disentangle the detrimental effect of aging and the brain’s adaptive response to such effect. Specifically, a more integrated network structure in the aging brain may reflect the deleterious effects of aging on brain function, which would be associated with poorer cognitive performance, or they may reflect compensatory response to age-related decline, which would be associated with better cognitive performance. Here, by using task-related fMRI and comparing functional brain networks within-subjects as a function of cognitive performance, we were able to directly link the increase in network integration in some brain areas, such as PFC, with age-specific cognition-enhancing effect. Our result is also consistent with a recent study comparing working memory task networks in OAs and YAs, where the task-related regions in OAs integrate more into the whole brain network when the task demands increased (Crowell et al. 2019). Nonetheless, it is still unclear whether increased integration in OAs serves as a generalizable mechanism of compensation in aging for brain systems supporting other cognitive functions besides memory. Also, given the more prevalent application of resting-state fMRI in neuroimaging studies, it remains an important question for future studies whether the age-related integration in task and resting state are mediated by shared or distinct underlying neural mechanisms.

### Age-related difference in connectivity reconfiguration

The second goal of the current study was to investigate whether OAs, relative to YAs, are more likely to reconfigure their patterns of PFC connections in a way that is associated with better memory performance. Accumulating evidence suggests that changes in functional connectivity are crucial for supporting cognitive functioning across varying task domains (Cole et al. 2013; Gallen et al. 2016) and demands (Hearne et al. 2017; Davis et al. 2018). Such changes imply selective recruitment and/or disengagement of regions with regard to the specific functional roles of the regions, the consequence of which would be a qualitative reconfiguration in the complex pattern of global connectivity (Hearne et al. 2017) rather than quantitative shifts (i.e., overall increase or decrease) of connectivity strength. Thus, in the current study, a correlation-based method was used to measure reconfiguration between high and low memory conditions, as this measure is sensitive to qualitative changes in connectivity patterns while concomitantly remaining insensitive to changes in simple connectivity features such as scaling or baseline shifts.

Reconfiguration can be assessed at the levels of whole-brain network (Cole et al. 2014; Gallen et al. 2016; Hearne et al. 2017), task-related subnetwork (Davis et al. 2018), or single regions (Geib, Stanley, Dennis, et al. 2017; Geib, Stanley, Wing, et al. 2017). In the current study, examining the reconfigurations in an ROI-wise manner gave us the best resolution to test our hypothesis regarding how the ‘hot spots’ of reconfiguration shift in relation to age. As expected, OAs showed greater-than-average reconfiguration in PFC regions, further supporting the possibility that the PFC plays a critical role in OAs during successful memory retrieval. Our results accord with a previous study on cognitive control, in which participants performed 64 different tasks during fMRI scan (Cole et al. 2013). In this study, brain regions in the fronto-parietal network – regions key to the cognitive control functions – exhibited the highest reconfiguration of connectivity between different tasks, and the level of reconfiguration between two tasks was driven by the dissimilarity of the two tasks. Our study confirmed the idea that reconfiguration of functional connectivity could reveal the functional importance of a brain region in driving the brain from one cognitive state to another (in our case, the high vs. low memory state). Our results further indicate that, the inter-subject variability of dynamic functional network reconfiguration may possibly be mediated by many factors such as age, which is an area largely unexplored.

### PFC compensates for MTL deficit by reconfiguration

Functional compensation implies a deficit for which the PFC is compensating. Here, an MTL deficit in OAs is suggested by low MTL reconfiguration **(Fig. 6B)** accompanied by less MTL integration with the rest of the brain network **(Figs. 3, 7C)**. These observations are in line with several previous studies, which have identified age-related reductions in MTL activity during memory retrieval (e.g., Daselaar, Fleck, Dobbins, et al. 2006; Dennis, Daselaar, et al. 2007; Dennis, Kim, et al. 2007; Davis et al. 2008; St. Jacques et al. 2012; Dew et al. 2012), pointing to an MTL deficit in healthy OAs. The presence of PFC compensation is indicated by the negative relationship between PFC and MTL connectivity reconfiguration **(Figs. 7A, 7B)**, which is consistent with a previous study showing a negative relationship between PFC and MTL activation in a memory encoding task in OAs (Hedden et al. 2005). Our results further extend these previous findings, showing an increased task-dependency of PFC connectivity profile when the capacity of MTL – the core regions of memory retrieval – is compromised by aging. These results may further the understanding of the way age-related compensation is systemically implemented in the brain.

It is worth noting that both OAs and YAs showed a negative correlation between memory-related PFC and MTL reconfiguration. This finding is not inconsistent with the idea that PFC reconfiguration compensates for MTL reconfiguration because compensatory mechanisms in OAs often exist also in YAs. It is commonly assumed that during episodic memory retrieval, MTL mediates the recovery of stored memory representations, whereas PFC mediates control processes during retrieval (e.g., monitoring the retrieval output) (Moscovitch and Melo 1997; Curran et al. 1997; Cabeza and Dennis 2012). Thus, in general, when MTL-mediated recovery is low, PFC-mediated control demands tend to increase. For example, MTL activity is greater for higher relative to low confidence hits (Daselaar, Fleck, and Cabeza 2006; Kim and Cabeza 2007), whereas PFC activity is greater for low than high confidence hits (Henson et al. 1999; Fleck et al. 2006; Kim and Cabeza 2007). This “seesaw” relationship between MTL and PFC is consistent with the negative correlation between memory-related reconfiguration in both YAs and OAs. Thus, rather than using a completely new mechanism, OAs are likely to be tapping a mechanism that also exists in YAs to compensate when the MTL output is low.

The shifts of reconfiguration in MTL and PFC probably indicate age-related shifts in cognitive architecture (Spreng and Turner 2019), although here it is unclear what specific retrieval processes are driving PFC reconfiguration in OAs. In the retrieval task investigated, participants must process a name of a visual scene (e.g., harbor), generate the visual image of an encoded scene that matches the name, and rate the quality of the retrieved scene. Thus, age-related differences may involve one or more of the three following three processes: (1) schema processing, (2) generation, and (3) post-retrieval monitoring. First, schema processing (e.g., how a harbor looks like) has been strongly associated with PFC regions, especially ventromedial PFC (van Kesteren et al. 2010; Preston and Eichenbaum 2013; Ghosh et al. 2014; Hebscher and Gilboa 2016; Gilboa and Moscovitch 2017), and there is evidence that OAs differentially rely on schematic knowledge compared to YAs (Mather et al. 1999; Spreng and Turner 2019). Second, generation is essential for recall tests, which are more dependent on PFC (Cabeza et al. 1997) than generation-independent item recognition tests. Due to weaker MTL-mediated memory recovery, OAs may have generated more alternative scenes from memory trying to match the verbal label. Finally, post-retrieval monitoring refers to assessing the validity of retrieved memory and is strongly linked with dorsolateral PFC regions (Henson et al. 1999; Fleck et al. 2006). In OAs, the reduction in memory quality may lead to increases in PFC-mediated ‘quality control’ (Jacoby, Shimizu, Velanova, et al. 2005; Jacoby, Shimizu, Daniels, et al. 2005; Dew et al. 2012; Spreng and Turner 2019). In short, the current study hints at potential systematic interplay between aging, large-scale brain network dynamics, and shifts in cognitive strategies, which may be further investigated in the future.

### Limitations

Considerable debate persists over the choice of atlas for defining nodes in network analyses (see Stanley et al., 2013). Atlas selection for node specification should depend on the particular questions being addressed by the experimenters. The AAL atlas is currently the most widely used atlas and has been used in multiple different network analyses of task-related fMRI data that are relevant to our current study (Gallen et al. 2016; Geib, Stanley, Wing, et al. 2017; Geib, Stanley, Dennis, et al. 2017). Therefore, using the AAL atlas can facilitate comparison with relevant works. The main methodological concern of AAL is that its larger ROIs may contain multiple functional regions, an issue that may be overcome by the use of functionally defined atlases (Yeo et al. 2011; Gordon et al. 2014; Power et al. 2014). However, most functionally defined atlases are derived from samples of younger adults, which could introduce bias in the current study where younger and older adults have been compared.

Modularity analysis on a common network enabled us to directly examine compensation-related hypotheses for specific sub-systems of interests in the brain. In our study, extra efforts were made to control for potential age-related biases in the common module assignment, and a supplementary analysis further supports the existence of the two modules of interest (PFC & MTL) in both age groups **(Fig. S1c)**. Still, we acknowledge that age-related difference in community structure may exist, as is indicated by recent studies (Geerligs et al., 2015; Gallen et al., 2016). It would be important to explore the common and distinct impact of aging on the community structure of functional network across different cognitive tasks. However, due to the limited sample size, it is beyond the scope of our current study to systematically characterize the age-related difference in the global network structure during memory retrieval, which remains an interesting question for future studies.

### Conclusions

Developing a better understanding of how some OAs functionally compensate to preserve high cognitive functioning remains an important goal in the research of aging and cognition, as this information may lead to potential interventions aimed at slowing cognitive decline. Our results are largely consistent with the previous studies identifying PFC compensation in aging, and yet develop a more detailed picture of how network interactions in the PFC help mediate compensatory function in aging, and how these interactions are implemented through both an increase of functional communication with other brain systems and a dynamic reconfiguration of patterns of functional connections.

## Acknowledgements

LD, SD, and RC were supported by grant NIH-MH114253-01. EW and RC were additionally supported by grants NIH-AG058574 and AG019731. ZM and BG were supported by grants F31AG060691 and F31MH114454, respectively.

## Footnotes

1 There has been considerable debate over which atlases should be used to define nodes for network analyses (Stanley et al., 2013). We contend that there is currently no “best” atlas available for use in network analyses, and that the selection of atlas for node specification should depend on the particular questions being addressed by the experimenters. We elected to use the AAL atlas because it is the most widely used atlas for defining nodes more generally, and because it has been used in multiple different network analyses of task-related fMRI data that are relevant to our current study (Gallen et al. 2016; Geib, Stanley, Dennis, et al. 2017; Geib, Stanley, Wing, et al. 2017). Consequently, comparison with relevant recent work is more straightforward and informative.

2 https://www.nitrc.org/projects/artifact_detect, developed by Shay Mozes and Susan Whitfield-Gabrieli, based on a previous tool by Paul Mazaika, Susan Whitfield, and Jeffrey C. Cooper.

